# Brain Connectivity and Behavioral Changes in a Spaceflight Analog Environment with Elevated CO_2_

**DOI:** 10.1101/2020.09.28.317404

**Authors:** Heather R. McGregor, Jessica K. Lee, Edwin R. Mulder, Yiri E. De Dios, Nichole E. Beltran, Igor S. Kofman, Jacob J. Bloomberg, Ajitkumar P. Mulavara, Rachael D. Seidler

**Affiliations:** Department of Applied Physiology and Kinesiology, University of Florida, Gainesville, FL; Institute of Aerospace Medicine, German Aerospace Center, Cologne, Germany; KBR, Houston, TX; NASA Johnson Space Center, Houston, TX

**Keywords:** Bed Rest, Spaceflight, CO_2_, Resting-state fMRI, Functional Connectivity, Cognition, Sensorimotor

## Abstract

Astronauts are exposed to microgravity and elevated CO_2_ levels onboard the International Space Station. Little is known about how microgravity and elevated CO_2_ combine to affect the brain and sensorimotor performance during and after spaceflight. Here we examined changes in resting-state functional connectivity (FC) and sensorimotor behavior associated with a spaceflight analog environment. Participants underwent 30 days of strict 6^°^ head-down tilt bed rest with elevated ambient CO_2_ (HDBR+CO_2_). Resting-state functional magnetic resonance imaging and sensorimotor assessments were collected 13 and 7 days prior to bed rest, on days 7 and 29 of bed rest, and 0, 5, 12, and 13 days following bed rest. We assessed the time course of FC changes from before, during, to after HDBR+CO_2_. We then compared the observed connectivity changes with those of a HDBR control group, which underwent HDBR in standard ambient air. Moreover, we assessed associations between post-HDBR+CO_2_ FC changes and alterations in sensorimotor performance. HDBR+CO_2_ was associated with significant changes in functional connectivity between vestibular, visual, somatosensory and motor brain areas. Several of these sensory and motor regions showed post-HDBR+CO_2_ FC changes that were significantly associated with alterations in sensorimotor performance. We propose that these FC changes reflect multisensory reweighting associated with adaptation to the HDBR+CO_2_ microgravity analog environment. This knowledge will further improve HDBR as a model of microgravity exposure and contribute to our knowledge of brain and performance changes during and after spaceflight.

## INTRODUCTION

Spaceflight elicits sensorimotor adaptive processes due to altered sensory inputs. Upon initial exposure to microgravity, crewmembers typically experience sensorimotor disturbances and disorientation, but they gradually adapt to body unloading, alterations in vestibular, and somatosensory inputs (Paloski et al., 1994, 1992; Reschke et al., 1998), as well as to headward fluid shifts (Lee et al., 2019; Roberts et al., 2017). In addition to microgravity, astronauts must also adapt to the mildly hypercapnic environment of the International Space Station (ISS) where carbon dioxide (CO_2_) levels average 0.5% -- approximately ten times higher than typical ambient levels (∼0.04%) on Earth (Law et al., 2014). On Earth, exposure to 3% CO_2_ over several hours produces headaches while resting. During mild exertion, subjects begin to report headaches at 2% CO_2_ (Schlute, 1964). Mild impairments in visuo-motor performance have been reported at levels as low as 1.2% CO_2_ (Manzey and Lorenz, 1998). Onboard the ISS, astronauts report CO_2_-related symptoms at lower CO_2_ levels than would be expected on Earth; astronauts are more prone to headaches (Law et al., 2014) and experience “space fog” characterized by slowed mental abilities and poor concentration (Kanas and Manzey, 2008; Welch et al., 2009). This suggests that microgravity and elevated CO_2_ levels may interact to negatively impact crewmember health and mission success.

Astronauts return to Earth in a microgravity-adapted state, exhibiting pronounced impairments in balance (Cohen et al., 2012; Paloski et al., 1994), locomotion (Bloomberg and Mulavara, 2003; Courtine and Pozzo, 2004; McDonald et al., 1996; Mulavara et al., 2018, 2010), and manual motor control (Manzey et al., 2000; Moore et al., 2019). Readaptation to Earth’s 1g environment takes place over the course of days or weeks (Miller et al., 2010; Mulavara et al., 2018, 2010).

Although the behavioral effects of spaceflight have been well-documented, the neural mechanisms underlying these changes remain largely unknown. We know even less about how exposure to microgravity and chronically elevated CO_2_ levels combine to affect the brain, cognition, and sensorimotor performance. Here, we investigated this issue using a well-established spaceflight analog, ‘head-down tilt bed rest’ (HDBR), combined with elevated levels of CO_2_ to better simulate environmental conditions (i.e., microgravity and mild hypercapnia) experienced during spaceflight onboard the ISS.

HDBR studies have become the predominant model for simulating microgravity on Earth, and studying its effects on the human body (Hargens and Vico, 2016; Pavy-Le Traon et al., 2007). In an HDBR study, healthy participants continuously lay in bed with their head tilted below the level of the feet. HDBR mimics some of the effects of microgravity exposure including axial body unloading, reduced somatosensory inputs (Reschke et al., 2009), deconditioning (Hargens and Vico, 2016), headward fluid shifts and upward shift of the brain within the skull (Koppelmans et al., 2017b), sensorimotor behavioral dysfunction (Koppelmans et al., 2017a; Reschke et al., 2009), and changes in brain activity (Cassady et al., 2016; Liao et al., 2015, 2012; Yuan et al., 2018a, 2018b, 2016; Zhou et al., 2014). In contrast to previous bed rest studies, here we used “strict” HDBR. To keep the head tilted down at a 6° angle, subjects were not permitted to use a standard pillow.

In the current study, we used resting-state functional magnetic resonance imaging (rsfMRI) to assess how HDBR+CO_2_ affects the brain’s large-scale functional network architecture. The brain has the highest metabolic activity of any organ in the human body, and the majority of its energy is consumed while in a state of wakeful rest (Lord et al., 2013; Raichle, 2010a, 2010b). While at rest, several networks of distributed brain regions exhibit spontaneous, low frequency BOLD signal fluctuations. Resting-state fMRI is a method used to estimate correlations within these resting-state networks. Resting-state network topographies closely correspond to structural connectivity and therefore correlations within these networks are thought to reflect functional connectivity (FC) (Biswal et al., 1995; Fox et al., 2005).

Demertzi and colleagues (2016) investigated resting-state FC changes in a single cosmonaut following a 169-day mission on the ISS. They reported reduced intrinsic connectivity within the right insula and ventral posterior cingulate cortex as well as reduced FC between the cerebellum and motor cortex relative to pre-flight measures (Demertzi et al., 2016). HDBR studies simulating microgravity with larger sample sizes have examined how simulated microgravity in ambient air have reported resting-state functional connectivity increases involving somatomotor and vestibular cortices, the cerebellum, insular and cingulate cortices, parietal regions, and the thalamus (Cassady et al., 2016; Liao et al., 2015, 2013, 2012; Zeng et al., 2016; Zhou et al., 2014). Connectivity increases may reflect increases in sensory weighting in microgravity or during HDBR, whereas connectivity decreases may be associated with reductions in use or reduced sensory weighting.

We have previously conducted a 70-day HDBR study in ambient air using a similar design, and an identical battery of sensorimotor and cognitive assessments to those employed here (Koppelmans et al., 2015). In our previous work, we have examined resting-state FC changes during HDBR in ambient air, and their associations with performance changes on identical behavioral assessments (Cassady et al, 2016). Methodology of the previous ambient air HDBR study differs with respect to the current HDBR+CO_2_ study. First, the ambient air HDBR study did not employ strict bed rest; subjects were permitted to use a standard pillow (which altered the degree of head tilt) and subjects were permitted to raise their head while eating. Second, under NASA’s directions, the previous ambient air HDBR study employed a different bed rest phase duration and a different testing timeline (Cassady et al, 2016). In the current study, we used similar analyses and identical regions of interests as Cassady et al (2016) to aid in drawing parallels between our results.

Lee et al (2019) contrasted the behavioral effects of HDBR+CO_2_ compared to our previous ambient air HDBR study. In light of the differing bed rest phase durations, comparisons relied on the rates (slopes) of performance change over time. Lee et al. (2019) reported differential pre-to post-bed rest performance changes on three assessments: 1) response variability on a rod and frame test, 2) time to complete a digit symbol substitution test, and 3) time to complete the Functional Mobility Test. Here, since we were interested in the additive effects of elevated CO_2_, we examined brain-behavioral associations for those behavioral measures that showed significantly different changes following HDBR+CO_2_ compared to HDBR in ambient air. As published by Lee et al (2019), the HDBR+CO_2_ group’s performance on the digit symbol substitution task followed a learning curve throughout the study, providing clear evidence of practice effects but no discernable effect of the bed rest intervention. For this reason, we have omitted data from the digit symbol substitution test from our brain-behavioral analyses.

The aim of the current study was to investigate brain and behavioral effects of prolonged exposure to simulated microgravity and elevated CO_2_ levels as a model of long-duration spaceflight onboard the International Space Station. Eleven subjects underwent 30 days of 6^°^ HDBR in 0.5% CO_2_. Cognitive and sensorimotor tests were administered before, during, and after the HDBR+CO_2_ intervention. Using both hypothesis-driven and hypothesis-free approaches, we examined: 1) time courses of resting-state FC changes from pre-, during-, to post-HDBR+CO_2_, and 2) relationships between FC changes and behavioral changes associated with HDBR+CO_2_. In light of previous work, we hypothesized that our 30-day HDBR+CO_2_ intervention would alter resting-state FC involving sensorimotor brain areas, regions involved in multisensory integration, and brain areas within the default mode network (Cassady et al., 2016; Demertzi et al., 2016; Koppelmans et al., 2017a; Liao et al., 2015, 2012; Zhou et al., 2014). We also investigated FC changes involving visual brain areas as elevated CO_2_ levels are hypothesized to contribute to the development of ophthalmic abnormalities and visual impairments in astronauts following long-duration spaceflight (Zwart et al., 2017). To determine the specificity of our findings, we conducted follow-up tests in which we compared the patterns of FC changes observed in the HDBR+CO_2_ group to those of a HDBR control group, which underwent strict HDBR in standard ambient air.

## METHODS

### Participants

All data were collected at :envihab, the German Aerospace Center’s medical research facility in Cologne, Germany. Prior to enrolment, all subjects passed psychological screening and an Air Force Class III equivalent physical examination. Subjects received monetary compensation for their participation. Study protocols were approved by the ethics commission of the local medical association (Ärztekammer Nordrhein) and institutional review boards at NASA, The University of Florida (HDBR+CO_2_ and HDBR control). All subjects provided written informed consent at :envihab.

#### HDBR+CO_2_ group

Twelve participants originally enrolled in the HDBR+CO_2_ study. One participant withdrew from the study on the first day of bed rest. Eleven participants completed the HDBR+CO_2_ study (6 males, 5 females; 25.3-50.3 years of age at study enrolment).

#### HDBR control group

Eight participants were included in a HDBR control group (6 males, 2 females; 27-46 years of age at study enrolment). Data from the HDBR control data were collected as part of a subsequent study which employed a 60-day bed rest phase. HDBR control data were also collected at :envihab approximately one year following data collection for the HDBR+CO_2_ group.

### Experimental Design

This experiment was part of the VaPER (Vision Impairment and Intracranial Pressure and Psychological :envihab Research) bed rest campaign. Subjects resided at :envihab for 58 consecutive days, during which they participated in several experiments. Our experiment consisted of 3 phases: a 14-day baseline data collection (BDC) phase, 30 days of strict 6^°^ head-down tilt bed rest phase with elevated CO_2_ (HDBR+CO_2_), followed by a 14-day recovery (R) phase (see Figure 1).

**Figure 1.**
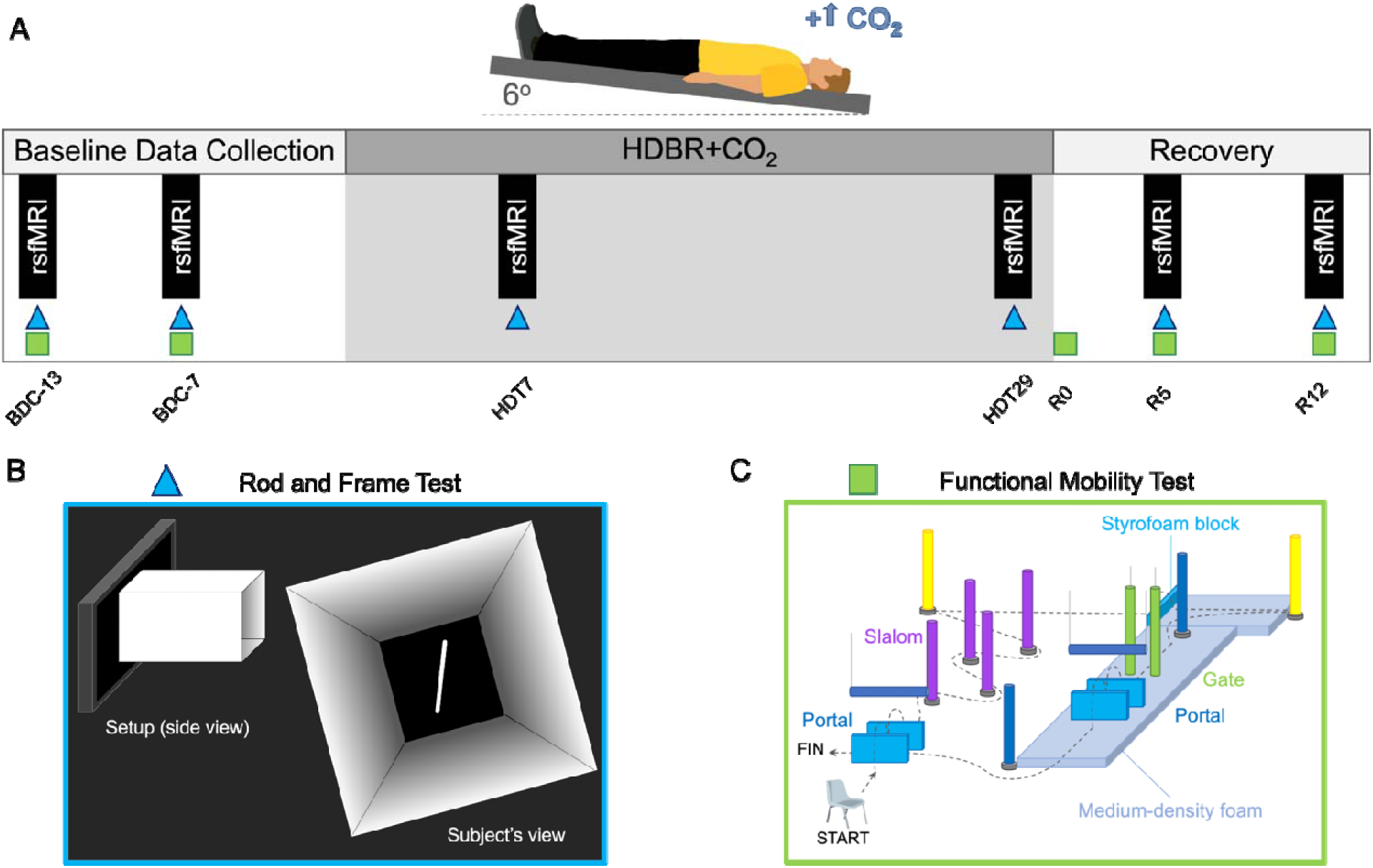
**A)** Subjects underwent a 15-day baseline data collection (BDC), 30 days of 6^°^ head-down tilt bed rest with elevated carbon dioxide (HDBR+CO_2_), followed by a 14-day recovery (R) phase. Resting-state fMRI scans occurred: 13 and 7 days before bed rest, on days 7 and 29 of the bed rest phase, and on the 6^th^ day (R5) and 13^th^ day (R12) during the recovery phase. **B)** Subjects performed a rod and frame test after each scan session. **C)** The Functional Mobility Test, an obstacle course, was administered twice during the baseline phase and three times following bed rest: on recovery days R0 (3 hours after first standing), R5, and R12.

#### Baseline Data Collection (BDC)

Subjects were admitted 15 days prior to bed rest for baseline data collection in standard ambient air (∼0.04% CO_2_). Subjects were ambulatory during the baseline phase, but were not permitted to leave the :envihab facility. Other than participating in study assessments, the subjects were not limited in their activities. Subjects slept in a horizonal (i.e., 0^°^ tilt) position. For both groups, baseline resting-state fMRI scans and behavioral assessments (described below) occurred 13 and 7 days prior to bed rest (BDC-13 and BDC-7, respectively). Behavioral tests were performed approximately 3 hours after neuroimaging sessions. Baseline behavioral assessments (except those requiring upright stance or locomotion) were performed while subjects laid in a 6^°^ HDT position.

#### Head-down Tilt Bed Rest

All subjects remained in the HDT position 24 hours a day. They were instructed to have at least one shoulder in contact with the mattress at all times, and were not permitted to raise their head or prop themselves up. To keep the head tilted down at a 6° angle, subjects were not permitted to use a traditional pillow. Subjects were permitted to use a 5 cm tall support for the head and neck only when laying on their side (Laurie et al., 2019; 2020). No pillows were used when subjects laid on their back. Subjects were also not permitted to raise, contract or stretch their legs during the bed rest phase.

#### HDBR+CO_2_ group

Subjects in the HDBR+CO_2_ group underwent 30 consecutive days of strict 6° HDBR in elevated CO_2_ (0.5%; 3.8 mmHg partial pressure of CO_2_). Resting-state fMRI scans and behavioral assessments occurred on days 5 and 29 of the HDBR+CO_2_ phase (HDT5 and HDT29, respectively). Behavioral assessments were conducted at the bedside approximately 3 hours after neuroimaging sessions while the subject was in the HDT position. The Functional Mobility Test, which required upright stance and locomotion, was not performed during the HDBR+CO_2_ phase.

#### HDBR control group

Subjects in the HDBR control group underwent 60 consecutive days of strict 6° HDBR in standard ambient air. Resting-state fMRI scans occurred on days 29 (HDT29) and 58 of the bed rest phase. Here we analyzed HDBR control group resting-state fMRI data for the HDT29 session only since both underwent a HDT29 scan session.

#### Recovery (R)

Following bed rest, all subjects underwent a 14-day recovery (i.e., day R0 to day R13) phase in standard ambient air (∼0.04% CO_2_). Subjects were ambulatory, were not restricted in their activities, and slept in a horizonal (i.e., 0^°^ tilt) position. Each subject received 1-hour rehabilitation sessions on days 3, 5, 7, 8, 9, and 11 of the recovery phase. Rehabilitation sessions involved active stretching and exercise regimens using a Bosu^®^ ball or coordination ladder to improve strength and coordination. For the HDBR+CO_2_ group, resting-state fMRI scans were acquired on R5 and R12. The the rod and frame test were performed after scan sessions on R5 and R12 while subjects laid in 6^°^ HDT position. The Functional Mobility Test was performed 3 times during the recovery phase: on days R0 (approximately 3 hours after first standing after bed rest), R5, and R12. The group differences in bed rest duration preclude direct comparisons between the two groups during the recovery phase. We therefore omitted recovery phase resting-state fMRI data for the HDBR control group.

As part of NASA’s standard measures assessments, blood draws were acquired 3 days prior to bed rest and on the first day after bed rest to calculate arterial partial pressure of carbon dioxide (PaCO_2_). We predicted that PaCO_2_ would increase from pre-to-post HDBR+CO_2_. Changes in PaCO_2_ were assessed using a one-tailed paired samples t-test.

### Behavioral Assessments

#### Rod & Frame Test

The rod and frame test assesses the extent to which subjects rely on a visual frame of reference for their perception of true vertical. As shown in Figure 1 (lower left), a 60-cm-long white tunnel-like frame was affixed to a flat screen computer monitor. A white rod was displayed on the screen inside the square frame. Subjects laid on their side at the edge of the bed and the end of the frame was positioned around their face to occlude any visual cues from the surrounding room. Subjects were then presented with random tilts of the rod and ±18° frame tilts for each trial, and they used a handheld controller to align the rod to Earth vertical (i.e., 0^°^ tilt). Each response was scored based on the deviation, in degrees, between perceived and true vertical. The primary outcome presented here is response consistency which refers to the within-subject variability of scores for a given rod/frame tilt combination. Since higher values reflect greater variability, here we refer to this measure as response variability.

#### Functional Mobility Test

Subjects’ locomotor function was assessed using the Functional Mobility Test (Figure 1, lower right). This test requires subjects to navigate an obstacle course by performing whole-body movements similar to those required to exit a vehicle after long-duration spaceflight (Mulavara et al., 2010; Koppelmans et al. 2013). The first half of the Functional Mobility Test was built on a hard surface while the second half was built on a medium-density foam floor. The foam base was used to reduce the reliability of proprioceptive input (Mulavara et al., 2010) and increase reliance on vestibular cues (Jeka et al., 2004). This course was comprised of 1) foam pylons arranged in a slalom pattern, requiring quick changes in heading direction, 2) “portals” consisting of a horizontally-hung foam obstacle between 2 styrofoam blocks which required subjects to bend at the waist and balance on one foot while stepping over the Styrofoam blocks, 3) a Styrofoam block located at the transition point between the hard floor and foam base, requiring subjects to balance on one foot on the unstable foam while clearing the Styrofoam block, and 4) a “gate” consisting of 2 vertically-hung pylons through which subjects entered sideways (i.e., shoulder first). Subjects started in a seated position. They were instructed to walk quickly and safely through the course without touching any obstacles. Here we present the time to complete the first of 10 trials as this measure is most sensitive to mobility changes.

Since the aim of the current study was to investigate brain and behavioral effects of our HDBR+CO_2_ intervention, we examined behavioral changes from immediately pre-to post-HDBR+CO_2_, as this is where the behavioral changes are most evident.

### MRI Image Acquisition

During all MRI scan sessions, a foam wedge was used to maintain the subject’s body in the 6° HDT position, but the head was horizontal in the MRI coil. For subjects in the HDBR+CO_2_ group only, a mask and tank system was used to maintain the 0.5% CO_2_ level throughout scan sessions during the bed rest phase.

All neuroimaging data were acquired on a 3-Tesla Siemens Biograph mMR scanner. T1-weighted anatomical images were collected using a MPRAGE sequence with the following parameters: TR = 1900 ms, TE = 2.44 ms, flip angle = 9°, FOV = 250 x 250 mm, matrix = 512 x 512, voxel size = 0.5 x 0.5 mm, 192 sagittal slices, slice thickness = 1 mm, slice gap = 0.5 mm. Whole-brain functional data were acquired using a T2*-weighted EPI sequence with the following parameters: TR = 2500 ms, TE = 32 ms, flip angle = 90°, FOV = 192 x 192 mm, matrix = 64 x 64, voxel size = 3 x 3 x 3.5 mm, slice thickness = 3.5mm, 37 axial slices, duration = 7 minutes. During the resting-state scan, participants were instructed to remain awake with their eyes open, fixate a red circle, and not think about anything in particular.

### Image Preprocessing

Neuroimaging data analyses were performed using the Statistical Parametric Mapping 12 software (SPM12; www.fil.ion.ucl.ac.uk/spm) and the CONN toolbox version 18a (Whitfield-Gabrieli and Nieto-Castanon, 2012) implemented in Matlab 2018b. Image preprocessing was identical for the HDBR+CO_2_ group and HDBR control group. Functional image preprocessing included spatial alignment to the first volume, slice timing correction, coregistration, and normalization with resampling to 2 mm isotropic voxels. Structural images were segmented into gray matter, white matter, and cerebrospinal fluid (CSF) tissue maps, and normalized to Montreal Neurological Institute (MNI) standard space.

To improve cerebellar normalization, we used CERES to segment the cerebellum from each subject’s anatomical image (Romero et al., 2017). CERES is an automated pipeline that uses a segmentation method called Optimized PatchMatch Label fusion (for details see Romero et al., 2017). CERES has been shown to yield more accurate cerebellar segmentations compared to other tools including the SUIT toolbox (Carass et al., 2018). CERES was used to segment the cerebellum from each native space anatomical scan. Generated cerebellar masks were used to extract the cerebellum from each subject’s session-wise T1 and coregistered functional images. The cerebellums extracted from functional and structural images were then normalized to the SUIT space template. The SUIT template was used because it retains more anatomical detail within the cerebellum compared to whole brain MNI templates, resulting in improved normalization for functional MRI (Diedrichsen, 2006).

Whole-brain functional images were spatially smoothed using a 5 mm full width at half maximum (FWHM) Gaussian kernel. The cerebellums extracted from functional scans were spatially smoothed using a 2 mm FWHM Gaussian kernel. A smaller smoothing kernel was used for the cerebellums so as to reduce smoothing across lobules and tissue types.

### Image Denoising

Since measures of resting-state FC are known to be influenced by head motion (Van Dijk et al., 2012), we used the ARTifact detection tool (ART; http://www.nitrc.org/projects/artifact_detect) to detect and reject motion artifacts. Within each resting-state run, a volume was deemed an outlier if the subject’s composite movement exceeded 0.9 mm or if the global mean intensity of the volume exceeded 5 standard deviations from the mean image intensity of the run. This corresponds to the intermediate default motion thresholds in the CONN toolbox (Whitfield-Gabrieli and Nieto-Castanon, 2012). ART generated a “scrubbing” regressor which identified outlier volumes. For the HDBR+CO_2_ group, the following number of volumes (± SD, range) were detected as outliers at each of the time points: BDC-13: 2.4 ± 3.3 SD, range=0-9; BDC-7: 1.5 ± 1.8 SD, range=0-4; HDT7: 0.36 ± 1.2 SD, range=0-4; HDT29: 0 outlier volumes; R5: 0.55 ± 1.8 SD, range=0-6; R12: 0.63 ± 1.3 SD, range=0-4. As 164 volumes were acquired during each resting-state run, less than 6% of the original data points (i.e., 9/164 volumes) were identified as outliers and regressed out across subjects and sessions. A repeated measures ANOVA followed by Bonferroni post-hoc tests revealed no statistically significant main effect of session on the number of outlier volumes [Wilks’ F(5,6)=1.426, p=0.336] for the HDBR+CO_2_ group. Including subject-wise regressors modelling the number of outlier volumes for each resting-state fMRI run resulted in qualitatively similar results. Similarly, for the HDBR control group, a repeated measures ANOVA followed by Bonferroni post-hoc tests revealed no significant main effect of session on the number of outlier volumes [Wilks’ F(2,6)=0.741, p=0.516].

Resting-state fMRI data are also influenced by noise arising from physiological fluctuations, primarily cardiac pulsations and respiration, which can exhibit spatial and spectral overlap with resting-state networks (Cole et al., 2010). We used the anatomical component-based noise correction (aCompCor) method to denoise our data (Behzadi et al., 2007). This method uses a principal components analysis approach to estimate noise signals in the white matter and CSF. For each session, the participant’s anatomical image was segmented into white matter and CSF masks. Resulting masks were eroded by one voxel to reduce partial volume effects, and were then applied to unsmoothed resting-state functional images. Unsmoothed functional images were used to avoid smoothing the BOLD signal across neighboring brain areas and different tissue types. BOLD signals were then extracted from the white matter and CSF used as noise regions of interest (ROIs) in a principal components analysis. The resulting significant components modelled the influence of noise as a voxel-wise linear combination of multiple estimated sources. For each run, 5 principal components were extracted for each of the white matter and CSF noise ROIs, and were included as nuisance regressors in our first-level GLM.

Each preprocessed resting-state run was denoised by regressing out confounding effects of head motion (6 head motion parameters and their 6 first-order temporal derivatives), 5 principal components modelling estimated noise from the white matter, 5 principal components modelling estimated noise from the CSF, and a “scrubbing” regressor modelling outlier volumes within the run. The resulting residuals were band-pass filtered between 0.008–0.09 Hz (Biswal et al., 1995; Damoiseaux et al., 2006) prior to being used in subject-level analyses.

### Subject-level analyses

We used two approaches to examine connectivity changes associated with HDBR+CO_2_: a hypothesis-based seed-to-voxel analysis and a hypothesis-free voxel-to-voxel analysis.

#### Seed-to-voxel analysis

For our seed-to-voxel analyses, we selected 13 *a priori* seed regions of interest (ROIs). We hypothesized that our HDBR+CO_2_ intervention would induce functional changes involving sensorimotor brain areas, regions involved in multisensory integration, and brain areas within the default mode network (Cassady et al., 2016; Demertzi et al., 2016; Koppelmans et al., 2017a; Liao et al., 2015, 2012; Zhou et al., 2014; Zeng et al., 2016). We used the same ROI coordinates as used in our previous resting-state fMRI study using an ambient air HDBR intervention (Cassady et al., 2016). Elevated CO_2_ levels are hypothesized to be one of the environmental factors contributing to the development of ophthalmic abnormalities and visual impairments in astronauts following long-duration spaceflight (Zwart et al., 2017). Therefore, we also included ROIs located within primary visual cortex and frontal eye field, the coordinates of which were determined using the automatic anatomic labeling (AAL) atlas included in the CONN toolbox. Table 1 shows coordinates for ROIs used in our seed-to-voxel analyses.

**Table 1.** Regions of interest (ROIs) and their coordinates used for seed-to-voxel analyses. L, left; R, right.

| Regions of Interest (ROIs) | MNI Coordinates |  |  |
| --- | --- | --- | --- |
|  | *x* | *y* | *z* |
| R premotor cortex (PMC) | 26 | -10 | 55 |
| R middle frontal gyrus (MFG) | 30 | -1 | 43 |
| L primary motor cortex (M1) | -38 | -26 | 50 |
| L putamen | -28 | 2 | 4 |
| L insular cortex | -41 | 12 | -9 |
| R vestibular cortex | 42 | -24 | 18 |
| L frontal eye field (FEF) | -27 | -9 | 64 |
| L posterior cingulate cortex (PCC) | -12 | -47 | 32 |
| R posterior parietal cortex (PPC) | 28 | -72 | 40 |
| R superior posterior fissure | 24 | -81 | -36 |
| L primary visual cortex (V1) | -10 | -70 | 10 |
| R cerebellar lobule V | 16 | -52 | -22 |
| R cerebellar lobule VIII | 20 | -57 | -53 |

ROIs included all voxels within a 4-mm-radius sphere centered on the ROI coordinates. We first extracted the time series from unsmoothed functional data within each ROI. For ROIs located in the cerebrum, ROI time series were drawn from the whole brain unsmoothed functional images (source image). For ROIs located in the cerebellum, ROI time series were drawn from the unsmoothed cerebellums extracted from functional images (source image). We then computed the spatial average of the BOLD time series across all voxels with the ROI.

We estimated FC between each ROI and the rest of the brain or within the cerebellum. For each ROI, we computed (bivariate) Pearson’s correlation coefficients between the mean ROI time series and the time series of every other voxel in the smoothed whole brain functional and within the cerebellum (i.e., smoothed cerebellums extracted from functional scans). Correlation coefficients were then Fisher Z-transformed to improve their normality. This procedure was performed for each of the subjects’ six resting-state scans separately.

#### Voxel-to-voxel analyses

We also used hypothesis-free voxel-to-voxel analyses. This allowed us to identify connectivity changes without restricting our analyses to our *a priori* selected ROIs. Here we examined changes in Intrinsic Connectivity Contrast (ICC), a measure which utilizes graph theory metrics to estimate global connectivity pattern strength (Martuzzi et al., 2011). This analysis involved computing (bivariate) Pearson’s correlation coefficients between the time course of each voxel with the time series of every other voxel in the brain (or within the cerebellum). The root mean square of each correlation was then computed. Correlation coefficients were then Fisher Z-transformed to improve their normality. This yielded one ICC map per run, each reflecting the connectivity strength (i.e., magnitude) at each voxel.

### Group-level Functional Connectivity Analyses

#### Time Course of Functional Connectivity Changes with HDBR+CO_2_

In this study, we were interested in the brain’s gradual adaptation to prolonged simulated microgravity and chronic CO_2_ elevation as opposed to the acute effects of elevated CO_2_ exposure. For both our seed-to-voxel and voxel-to-voxel analyses, we used the longitudinal model shown in Figure 2 to assess the time course of FC changes from pre-, during-, to post-bed rest. Our aim was to identify those brain regions that exhibited stable FC across the 2 baseline time points, then showed a gradual change in FC during HDBR+CO_2_, followed by a gradual return to baseline over the 2 post-bed rest time points. In this way, subjects served as their own controls.

**Figure 2.**
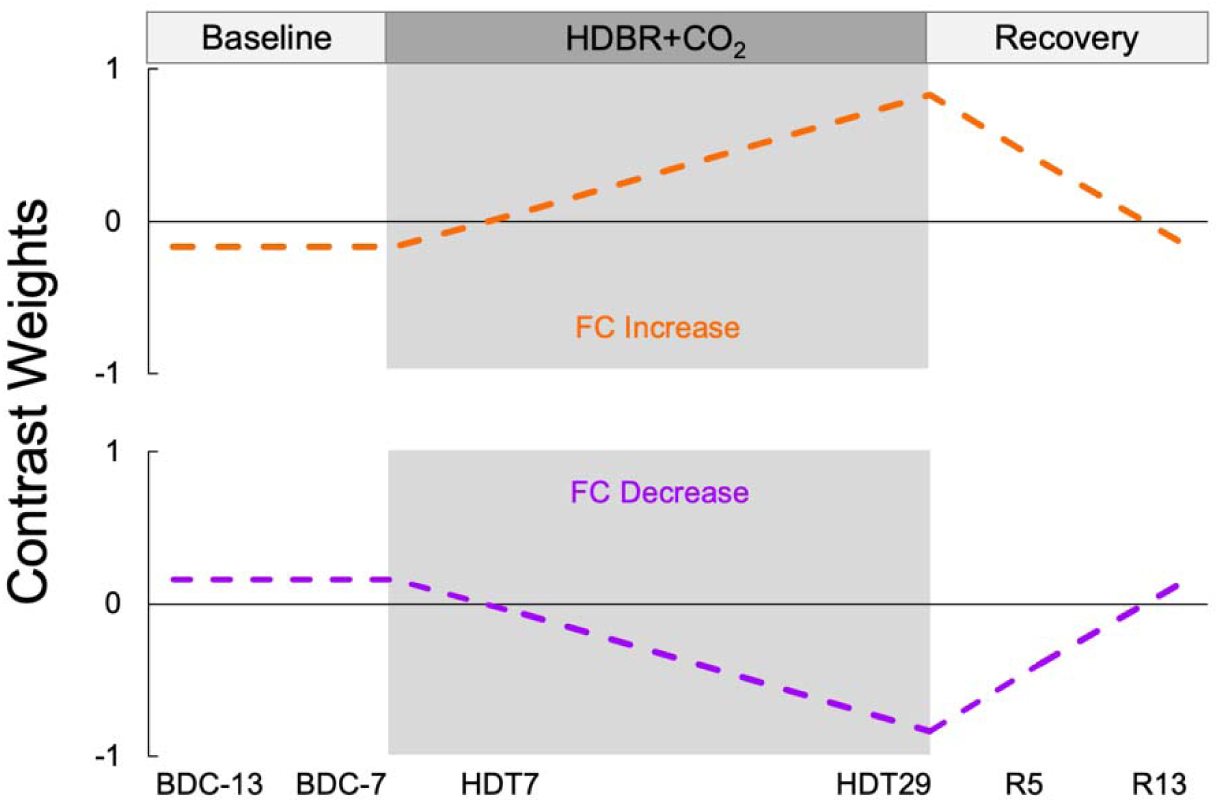
Contrast models used to assess time course of FC changes of the HDBR+CO_2_ group. Contrast weights used in the HDBR+CO_2_ group analyses. The gray shaded region indicates the HDBR+CO_2_ phase. The contrast in orange models a gradual FC increase during bed rest followed by a gradual FC decrease back to baseline level following bed rest. The contrast in purple models a gradual FC decrease during bed rest followed by a gradual FC increase back to baseline level following bed rest. BDC, baseline data collection; HDBR+CO_2_, head-down tilt bed rest with elevated carbon dioxide; R, recovery.

The HDBR+CO_2_ group and HDBR control group differed in bed rest duration (30 vs. 60 days) and data collection timeline (6 vs 5 sessions). This precluded direct comparison of the groups in our GLM. In light of this, we performed GLM analyses on the HDBR+CO_2_ group data only. Based on the results of these analyses, we performed follow-up assessments using resting-state fMRI data from the HDBR control group (detailed below).

For these analyses, our hypothesized model weights as a within-subjects contrast in our general linear model (GLM). The between-subjects regressor of interest was the grand mean. Age and sex were included as between-subjects covariates of no interest. Our analysis is analogous to that used by Cassady et al (2016). An uncorrected voxel threshold of *p* < 0.001 was set, and results were considered statistically significant at a cluster-level threshold of *p* < 0.05 (two-tailed), corrected for multiple comparisons according to the false discovery rate (FDR) method.

#### Time Course of Functional Connectivity Changes for HDBR control group

We performed follow-up assessments using the resting-state fMRI data from the HDBR control group. This was done to examine if the FC changes identified by our HDBR+CO_2_ longitudinal analyses were due to undergoing strict HDBR alone or due to the combination of strict bed rest and elevated CO_2_. We used HDBR control group resting state data from BDC-13, BDC-7, and HDT29 sessions as the both groups underwent scans on these days. For each significant cluster identified by our HDBR+CO_2_ longitudinal analysis, we generated a binary mask of the cluster. The cluster mask was used to extract FC or ICC values for each subject in the HDBR control group for sessions BDC-13, BDC-7, and HDT29. If a cluster resulted from a seed-to-voxel analysis, we extracted average FC values between the ROI and all voxels within the cluster mask. If a cluster resulted from a voxel-to-voxel analysis, we extracted average ICC values across all voxels within the cluster mask.

We then performed a split-plot ANOVA using IBM® SPSS® Statistics 21 to compare the time course of connectivity changes across BDC-13, BDC-7 and HDT29 between the HDBR+CO_2_ group and the HBDR control group. If the connectivity changes observed in the HDBR+CO_2_ group from BDC-13 to HDT29 are indeed due to the combination of strict HDBR and elevated CO_2_, it follows that the time courses of connectivity changes will differ between the two groups (i.e., a significant group x session interaction). However, if the connectivity changes observed in the HDBR+CO_2_ do not differ from those of the HDBR control group, then this would suggest that the connectivity change was a result of undergoing strict HDBR.

#### Behavioral Analyses

We have recently examined cognitive and motor performance alterations following 6^°^ HDBR+CO_2_ compared to 6^°^ HDBR with ambient air (Lee et al., 2019). These two bed rest studies employed an identical battery of behavioral tests (Koppelmans et al., 2015). Since we were interested in the additive effects of elevated CO_2_, here we examined only those behavioral measures that showed significantly different changes following HDBR+CO_2_ compared to ambient air HDBR. Differential performance changes were found for the following behavioral measures: i) response variability on the rod and frame test (referred to as response consistency in previous work), and ii) time to complete the full Functional Mobility Test.

Here we tested if pre-to post-HDBR+CO_2_ FC changes were associated with behavioral changes (i.e., a change-change correlation). For each of the behavioral measures above, we computed each subject’s change in behavior from the last baseline time point (BDC-7) to the end of bed rest (HDT29; R0 for the Functional Mobility Test only).

#### Associations Between Functional Connectivity and Behavior Changes

We then performed group-level GLM analyses to test if pre-to post-HDBR+CO_2_ FC changes were associated with behavioral changes. Here we examined FC from 7 days before bed rest (BDC-7) to day 29 of bed rest (HDT29). A separate analysis was performed for each behavioral test. For these analyses, we calculated the change in each subject’s performance from before bed rest to the end of bed rest. The regressor of interest for our between-subjects contrast modeled the performance changes across subjects. Subject age, sex, and the grand mean were included as covariates of no interest. Our within-subjects contrast modeled the change in FC from pre-to post-bed rest. These analyses allowed us to examine those areas of the brain in which FC changes following HDBR+CO_2_ were significantly associated with performance changes. An uncorrected voxel threshold was set at *p* < 0.001, and results were thresholded at a FDR-corrected significance level of p <0.05 (two-tailed, cluster-wise).

In total, 56 tests were conducted using GLMs (13 seed-to-voxel and 1 voxel-to-voxel analyses conducted for the time course analysis and each of the 3 brain-behavior associations). We used the Benjamini-Hochberg procedure to control the false discovery rate at 0.05 (Benjamini and Hochberg, 1995). Cluster-level uncorrected p value obtained by each GLM analysis were submitted to Benjamini and Hochberg’s FDR procedure with αFDR = 0.05 (Chumbley et al., 2010). A cluster was considered statistically significant if its uncorrected cluster size *p* < 0.00089 (i.e., (1/56) x 0.05).

## RESULTS

A paired samples t-test revealed a small but significant increase in PaCO_2_ from pre-(41.4 mmHg) to post-(43.4 mmHg) HDBR+CO_2_ (t(10)=2.16, p=0.028, one-tailed).

### Time Course of Functional Connectivity Changes with HDBR+CO_2_

#### Seed-to-voxel analyses

Seed-to-voxel analyses revealed that right vestibular cortex exhibited increases in FC with right cerebellar crus I and II during HDBR+CO_2_, followed by a reduction in connectivity following bed rest (Figure 3A). A split plot ANOVA examining group differences in longitudinal FC changes revealed a non-significant group x session interaction (F(2,34)=2.519, p=0.095) although there was a trend.

**Figure 3.**
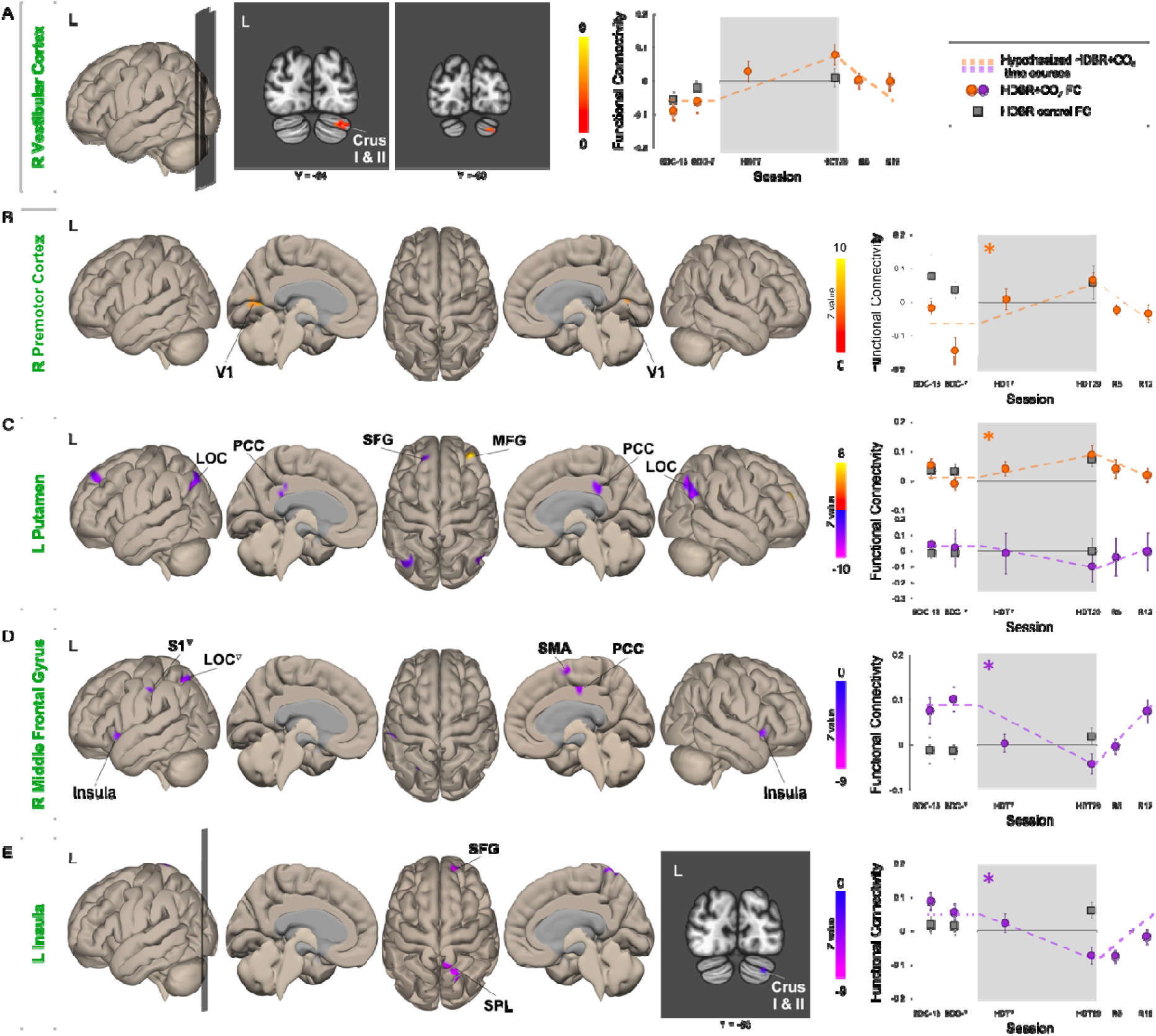
Time Course of Functional Connectivity Changes with HDBR+CO_2_. Results of our seed-to-voxel analyses, with each ROI indicated in green font. Significant clusters that exhibited FC increases during HDBR+CO_2_ followed by a post-bed rest reversal are shown in warm colors. Significant clusters that exhibited FC decreases during HDBR+CO_2_ followed by a post-bed rest reversal are shown in purple. Plots on the right represent session-wise functional connectivity values averaged across all clusters. Results from the HDBR+ CO_2_ group are indicated by colored circles. FC values across all clusters were extracted from the HDBR control group and are indicated by grey squares. ^▽^ indicates a cluster that did not survive the post hoc Benjamini-Hochberg FDR correction. * indicates a significant group x session interaction. Error bars represent standard error. L, left; R, right; FC, functional connectivity; BDC, baseline data collection; HDT, head-down tilt; R, recovery.

Our ROI in right premotor cortex exhibited FC increases with bilateral primary visual cortex (V1) during HDBR+CO_2_, which decreased after bed rest (Figure 3B). A split-plot ANOVA revealed a significant group x session interaction on FC values (F(2,34)=4.485, p=0.019). This indicated differential patterns of FC changes from the baseline phase to the HDT29 session in which the HDBR+CO_2_ group exhibited a greater increase during the bed rest phase.

The ROI in left putamen showed increased FC with clusters in right superior and middle frontal gyrus during HDBR+CO_2_, which reversed after bed rest (Figure 3C). The time course of FC for the HDBR+CO_2_ group was significantly different from that of the HDBR control (F(2,34)=4.049, p=0.026) such that the HDBR+CO_2_ showed greater FC increases during bed rest. The ROI in left putamen also exhibited FC decreases between left superior frontal gyrus, bilateral lateral occipital cortices, and bilateral posterior cingulate cortices during bed rest, followed by a reversal post-bed rest (Figure 3C). A split-plot ANOVA revealed a significant group x session interaction on FC values (F(2,34)=3.961, p=0.028), suggesting that FC decreases during bed rest were greater for the HDBR+CO_2_ group.

The ROI in right middle frontal gyrus exhibited FC decreases with a distributed network during bed rest including bilateral insular cortices, left primary somatosensory cortex, left lateral occipital cortex, right supplementary motor area, and right posterior cingulate cortex (Figure 3D). However, the clusters in left primary somatosensory cortex and left lateral occipital cortex did not survive post hoc Benjamini-Hochberg FDR correction. Consequently, FC values from these 2 clusters were omitted from the FC plot on the far right of Figure 3D and omitted from our split-plot ANOVA. The longitudinal pattern of FC changes significantly differed between the two groups (F(2,34)=3.92, p=0.029) such that the HDBR+CO_2_ showed greater FC decrease during bed rest.

Finally, left insula showed decreased FC with right SFG, right SPL, and cerebellar crus I and II, followed by a reversal after bed rest (Figure 3E). Cluster coordinates and statistics are presented in Table 2. A split-plot ANOVA revealed a significant group x session interaction on FC value (F(2,34)=11.046, p= 0.0002) demonstrating that the HDBR+CO_2_ group exhibited greater FC decreases from the baseline phase to HDT29 compared to the HDBR control group.

**Table 2.**
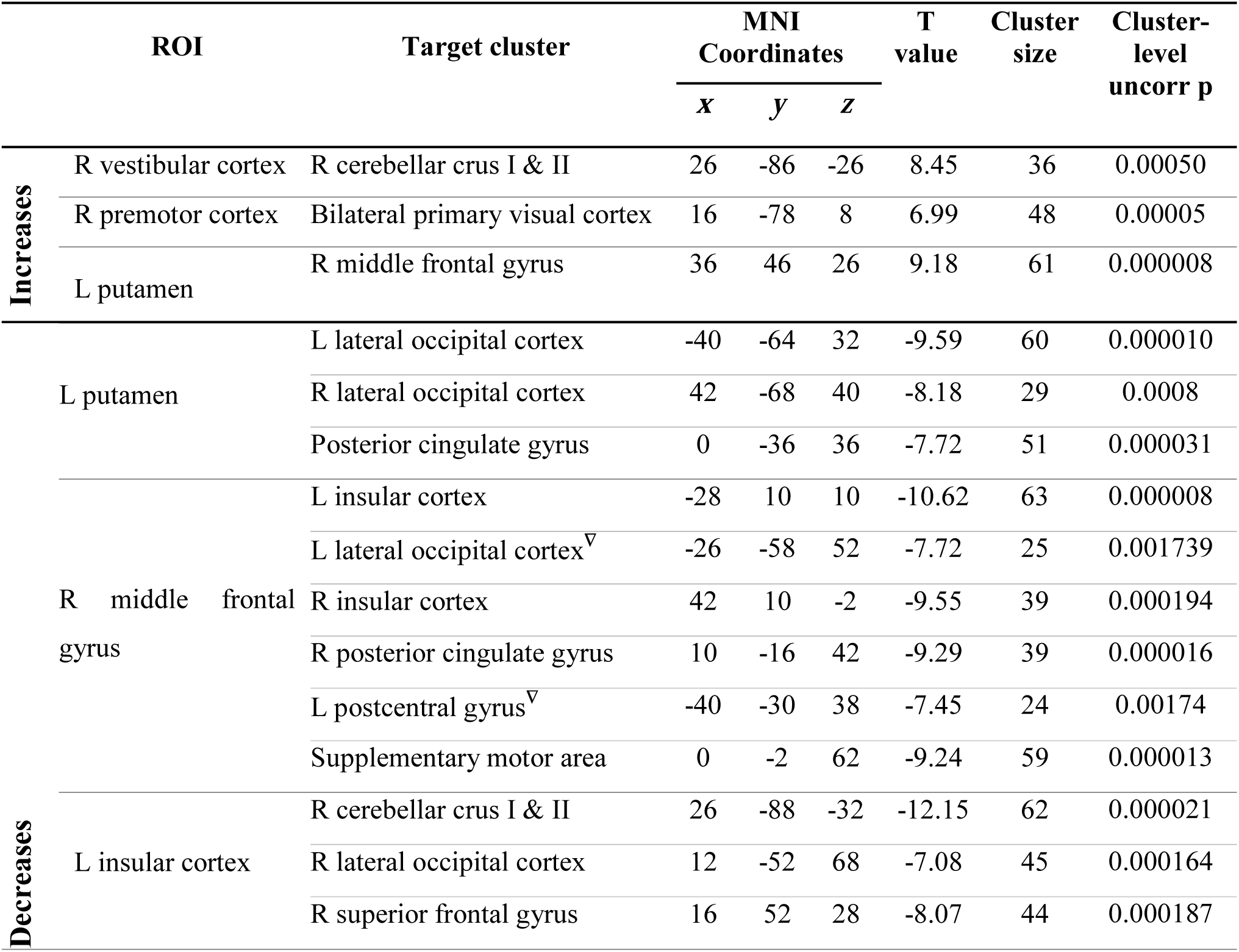
Significant clusters from our seed-to-voxel analyses assessing FC change time courses. Top rows indicate ROIs and clusters that exhibited FC increases during HDBR+CO_2_ followed by FC decrease during the post-bed rest recovery period. Bottom rows indicate ROIs and clusters that exhibited FC decreases during HDBR+CO_2_ then FC increase following bed rest. Cluster size refers to the number of voxels in each cluster. Cluster size uncorr p refers to uncorrected cluster size *p*-values. ^▽^ indicates a cluster that did not survive the post hoc Benjamini-Hochberg FDR correction. L, left; R, right; ROI, region of interest; uncorr, uncorrected.

#### Voxel-to-voxel analyses

Voxel-to-voxel analyses revealed a significant cluster within right primary visual cortex. Intrinsic connectivity contrast (ICC) within right V1 decreased during HDBR+CO_2_ then increased during the post-bed rest recovery phase (Figure 4, Table 3). A split-plot ANOVA examining group differences in longitudinal ICC changes revealed a non-significant group x session interaction (F(2,34)=2.172, p=0.130) although there was a trend.

**Table 3.** Significant cluster from our voxel-to-voxel analyses assessing FC change time courses. Cluster size refers to the number of voxels in each cluster. Cluster-level uncorr p refers to uncorrected cluster size *p*-values. R, right; uncorr, uncorrected.

| Cluster | MNI Coordinates |  |  | T value | Cluster size | Cluster-level uncorr p |
| --- | --- | --- | --- | --- | --- | --- |
|  | x | y | z |  |  |  |
| R primary visual cortex | 6 | -88 | 10 | -9.85 | 46 | 0.00001 |

**Figure 4.**
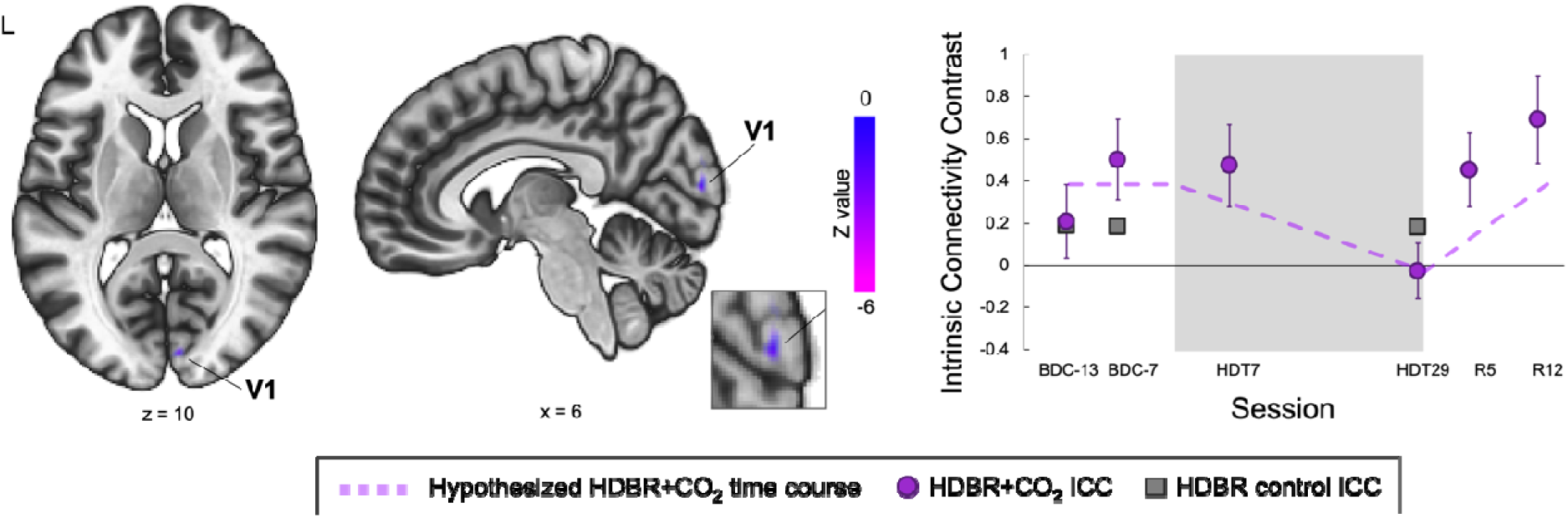
Time Course of Intrinsic Connectivity Contrast (ICC) with HDBR+CO_2_. Results of our voxel-to-voxel analyses. A cluster in right primary visual cortex exhibited decreased connectivity during HDBR+CO_2_ followed by a reversal during the recovery phase (colored circles). ICC values across all clusters were extracted from the HDBR control group and are indicated by grey squares. The inset shows an enlarged sagittal view of the cluster. The plot on the right reflects average Fisher Z-transformed intrinsic connectivity contrast (ICC) estimates across all subjects. Error bars represent standard error. Dashed lines represent our hypothesized longitudinal model. L, left; BDC, baseline data collection; HDT, head-down tilt; R, recovery.

### Associations Between Functional Connectivity and Behavior Changes

Lee and colleagues (2019) recently contrasted behavioral changes during our HDBR+CO_2_ intervention compared to our previous ambient air HDBR study. Here we examined brain-behavior associations for those behavioral measures that exhibited differential changes during HDBR+CO_2_ compared to our previous ambient air HDBR study. Lee et al. reported that response variability on the rod and frame test remained consistent during HDBR in ambient air, but decreased during HDBR+CO_2_. While both interventions resulted in slowed performance on the Functional Mobility Test, the degree of slowing was greater for the HDBR+CO_2_ group. Therefore, subjects who underwent the HDBR+CO_2_ intervention exhibited improved consistency of visual orientation perception compared to the ambient air HDBR group. However, the HDBR+CO_2_ group exhibited greater mobility impairments compared to the ambient air HDBR group (Lee et al., 2019). Figure 5 shows the evolution of each of the behavioral measures across all sessions for the HDBR+CO_2_ group only.

**Figure 5.**
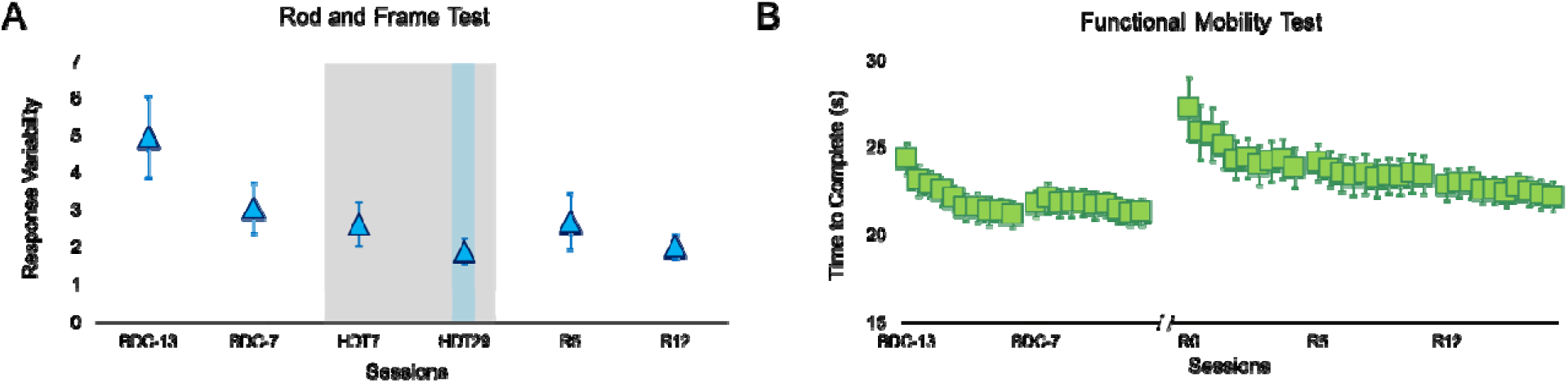
Performance on behavioral assessments. Session-wise group average of response variability on the rod and frame test **(A)** and time to complete each trial of the Functional Mobility Test **(B)**. Error bars represent SEM. The gray shaded region indicates the HDBR+CO_2_ phase. Colored bars indicate data used for pre-to post-bed rest performance comparisons. BDC, baseline data collection; HDT, head-down tilt; R, recovery.

We also examined how pre-to post-bed rest FC changes related to behavioral changes across subjects in the HDBR+CO_2_ group.

#### Rod and Frame Test

On average, subjects’ responses showed reduced variability (i.e., increased consistency) during HDBR+CO_2_ (Figure 5A). However, as shown in scatterplots in Figure 6, individual subjects differed in the direction and extent of response variability changes with HDBR+CO_2_. We found that post-bed rest FC decreases between left frontal eye field (FEF) and right lateral occipital cortex (LOC) were associated with decreased response variability (i.e., increased consistency) following bed rest (see Figure 6A). FC decreases between left insula and left primary somatosensory cortex (S1) were also associated with decreases in response variability. Finally, we also observed a relationship between connectivity changes within the default mode network and performance changes on the rod and frame test (Figure 6C). Post-bed rest FC decreases between left posterior cingulate cortex, left superior frontal gyrus, and right superior parietal lobule (SPL) were associated with decreases in response variability. The scatterplots on the right show the relationship between pre-to-post-HDBR+CO_2_ performance changes, which were used as the regressor of interest, and FC changes across subjects. Scatterplots are intended for illustrative purposes. A further correlation analysis was not performed because the nonindependence of such an analysis would inflate the effect size (Poldrack & Mumford, 2009). Cluster coordinates and statistics are presented in Table 4. Results were qualitatively similar if we used the subjects’ slope of behavioral change across BDC-7, HDT7, and HDT29 as our regressor of interest.

**Table 4.** MNI coordinates of clusters that showed post-bed rest FC changes that were significantly associated with changes in response variability on the rod and frame test. Cluster size refers to the number of voxels in each cluster. Cluster-level uncorr p refers to uncorrected cluster size *p*-values. ROI, region of interest; L, left; R, right; uncorr, uncorrected.

| ROI | Target cluster | MNI Coordinates |  |  | T value | Cluster size | Cluster-level uncorr p |
| --- | --- | --- | --- | --- | --- | --- | --- |
|  |  | x | y | z |  |  |  |
| **L frontal eye field** | R lateral occipital cortex | 42 | -76 | 40 | 12.99 | 48 | 0.000029 |
| **L insular cortex** | L postcentral gyrus | -40 | -36 | 48 | 12.57 | 50 | 0.00028 |
| **L posterior cingulate cortex** | L superior frontal gyrus | -34 | 12 | 64 | 9.26 | 38 | 0.000123 |
|  | R superior parietal lobule | 6 | -54 | 36 | 9.37 | 35 | 0.000199 |

**Table 5.** MNI coordinates of clusters that showed post-bed rest FC changes that were significantly associated with changes in time to complete the Functional Mobility Test. Cluster size refers to the number of voxels in each cluster. Cluster-level uncorr p refers to uncorrected cluster size *p*-values. ROI, region of interest; L, left; R, right; uncorr, uncorrected.

| ROI | Target cluster | MNI Coordinates |  |  | T value | Cluster size | Cluster-level uncorr p |
| --- | --- | --- | --- | --- | --- | --- | --- |
|  |  | x | y | z |  |  |  |
| **R primary motor cortex** | L V5 | -44 | -76 | 2 | 12.79 | 34 | 0.000101 |
| **R premotor cortex** | L cerebellar crus I | -34 | -70 | -26 | 10.57 | 51 | 0.000002 |
| **L primary visual cortex** | R inferior parietal lobule | 44 | -34 | 24 | -10.38 | 25 | 0.000279 |

**Figure 6.**
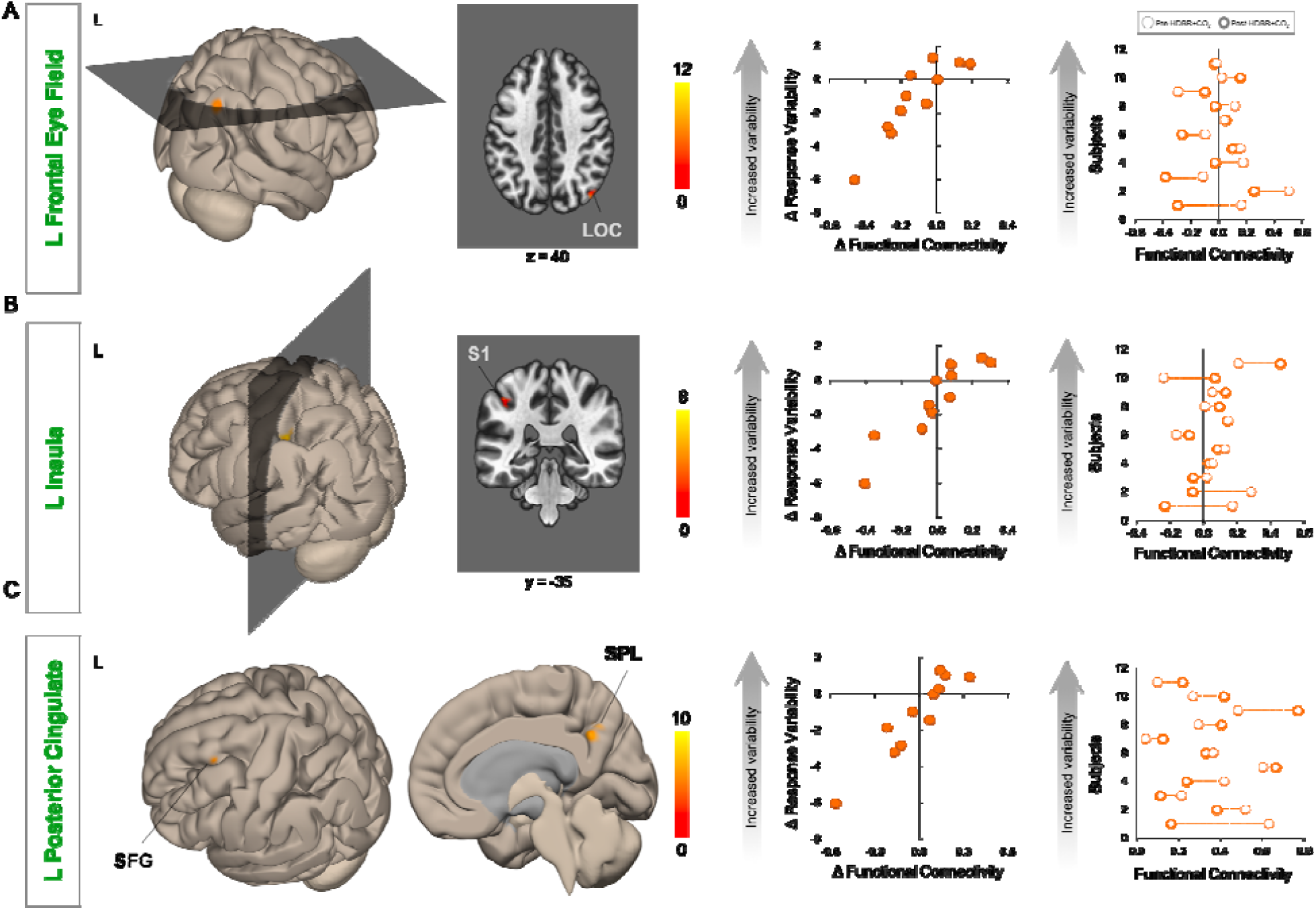
Pre-to Post-HDBR+CO_2_ changes in FC and response variability on the rod and frame test. Results of our seed-to-voxel analyses, with each ROI indicated in green font. Columns 1 and 2 show clusters that exhibited FC increases that were significantly associated with response variability increases from pre-to post-bed rest. Changes in response variability are plotted against functional connectivity changes in column 3 to illustrate the direction of the effect. The plot in column 4 shows subject-wise functional connectivity values before and after HDBR+CO_2_. Subjects are rank-ordered along the y-axis according to increasing response variability. L, left; LOC, latersal occipital cortex; S1, primary somatosensory cortex; SFG, superior frontal gyrus; SPL, superior parietal lobule; BDC, baseline data collection; HDT, head-down tilt; FC, functional connectivity; HDBR+CO_2_, head-down tilt bed rest with elevated CO_2_.

**Figure 7.**
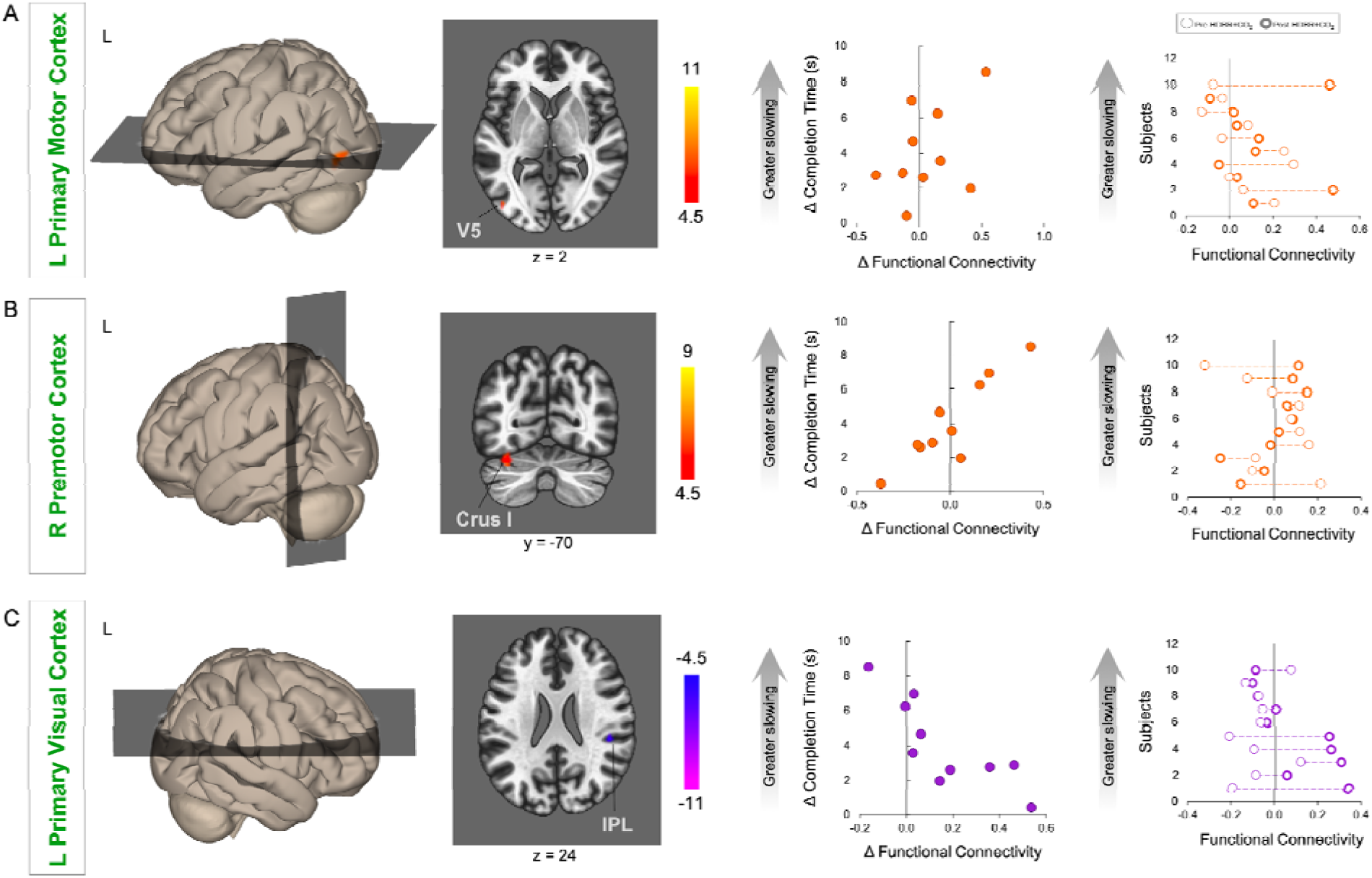
Pre- to Post-HDBR+CO_2_ changes in FC and time to complete the Functional Mobility Test. Results of our seed-to-voxel analyses, with each ROI indicated in green font. Clusters that exhibited post-HDBR+CO_2_ FC increases that were significantly associated with increases in Functional Mobility Test completion time. Changes in completion time are plotted against functional connectivity changes in column 3 to indicate the direction of the effect. The plot on the far right shows subject-wise functional connectivity values before and after HDBR+CO_2_. Subjects are rank-ordered along the y-axis according to increasing slowing of mobility. L, left; R, right; IPL, inferior parietal lobule.

#### Functional Mobility Test

All subjects exhibited slowing on the Functional Mobility Test (i.e., increases in completion times) following HDBR+CO_2_ (see Figures 5C, Figure 8), however, the extent of slowing varied across subjects. Post-bed rest FC increases between left primary motor cortex and left visual area V5 were associated with greater slowing on the Functional Mobility Test (Figure 8A). FC increases between right premotor cortex and left cerebellar crus I were also associated with greater slowing on the Functional Mobility Test following bed rest (see Figure 8B). Post-bed rest FC decreases between left primary visual cortex and right inferior parietal lobule were associated with greater slowing on the Functional Mobility Test. In contrast, subjects who exhibited increased FC between these areas showed less slowing. Cluster statistics are summarized in Table 6.

#### Voxel-to-voxel analyses

Voxel-to-voxel brain-behavioral analyses did not yield any statistically significant clusters.

## DISCUSSION

The aim of the current study was to investigate brain and behavioral effects of prolonged exposure to simulated microgravity and elevated CO_2_ levels as a model of long-duration spaceflight onboard the International Space Station. To this end, we tested how 30 days of head-down tilt bed rest in 0.5% CO_2_ affected resting-state functional brain connectivity. Our hypothesis-driven approach revealed that 30 days of HDBR in elevated CO_2_ resulted in resting-state FC increases among vestibular, motor, and primary visual brain areas, which decreased following bed rest. These brain areas are involved in balance and spatial orientation, the planning, control, coordination and adaptation of voluntary movement, and low-level visual processing, respectively. In contrast, we found decreases in resting-state FC involving motor, somatosensory, vestibular, cognitive, and multimodal integration regions during bed rest, which increased following bed rest. These are brain areas implicated in voluntary movement planning, sensorimotor control and somatosensation, balance and spatial orientation, attention and awareness (Leech and Sharp, 2014), and visuospatial perception, respectively. Our hypothesis-free analysis revealed that connectivity within right primary visual cortex gradually decreased during the HDBR+CO_2_ invention, then increased during the post-bed rest recovery phase. However, upon comparing the longitudinal ICC change of the HDBR+CO_2_ group with those of the HDBR control group, the two groups did not show a significantly different pattern of ICC change -- although there was a trend toward greater decreases for the HDBR+CO_2_ group.

Liao and colleagues (2012; 2013; 2015) have previously investigated resting-state activity changes during short duration HDBR interventions in ambient air. Employing a 7-day 6^°^ HDBR intervention, they examined changes in the amplitude of low frequency fluctuations (ALFF) at baseline and on each day of their HDBR intervention. During bed rest, subjects exhibited increased ALFF in anterior cingulate cortex and the posterior lobe of the cerebellum as well as ALFF decreases in posterior cingulate cortex and left paracentral lobule (Liao et al., 2015). Here, we similarly found FC decreases involving posterior cingulate cortex during HDBR+CO_2_. However, it is worth noting that Liao et al. (2015) did not leverage the longitudinal nature of their experimental design; they performed a one-way ANOVA across all time points followed by pairwise comparisons between each day of the HDT period versus the baseline phase.

Recent studies have examined the effects of long-duration HDBR in ambient air on large scale resting-state networks (Cassady et al., 2016; Zhou et al., 2014). Zhou et al (2014) used graph-theory based analyses to investigate pre-to post-bed rest FC changes. Following 45 days of HDBR in ambient air, subjects showed decreased degree centrality in the left anterior insula and dorsal anterior cingulate cortex. In the current study, we similarly observed widespread FC decreases involving insular cortices.

The current study built upon our recent work which demonstrated FC changes associated with 70 days of HDBR in ambient air. As in the current study, Cassady et al (2016) assessed connectivity changes from pre-, during, to post-bed rest using an analysis which incorporated both between- and within-subjects factors in the analysis. This type of analysis leveraged the longitudinal nature of the experimental design. It allowed for the identification of not only pre-to post-bed rest changes, but also the time course of FC changes throughout bed rest, using subjects’ baseline measures as their own controls. Cassady et al (2016) showed gradual FC increases between primary motor and somatosensory cortices, and between vestibular cortex and lobule VI within the ipsilateral cerebellar hemisphere, which decreased after bed rest. They also showed gradual FC decreases between posterior parietal cortex and temporal-occipital cortex, between cerebellar lobule VIII and postcentral gyrus, and between the superior posterior fissure and crus I of the cerebellum. In the current study, we used similar analyses and identical regions of interests as Cassady et al (2016) to aid in drawing parallels between our results. Similar to the 70-day ambient air HDBR results, we observed gradual FC increases between vestibular cortex and the cerebellum with bed rest, albeit the cerebellar cluster here spanned Crus I and II of the contralateral hemisphere (which are part of the default mode network and networks involved in cognitive control (Buckner et al., 2011). Reinforcing this similarity with the results of Cassady et al. (2016), the HDBR control group included in the present study exhibited a similar FC increase between vestibular cortex and crus I and II of the cerebellum as the HDBR+CO_2_ group. This finding suggests that this result is driven by undergoing strict HDBR as opposed to the combination of HDBR and elevated CO_2_. A notable qualitative difference between the results of the current study and those of Cassady et al (2016) is that HDBR+CO_2_ resulted in a wider spread pattern of functional decoupling among motor, somatosensory, vestibular, cognitive, and multimodal integration brain areas.

To our knowledge, only one published study has examined resting-state FC changes following spaceflight. In their case study of a single cosmonaut who spent 169 days onboard the ISS, Demertzi et al (2016) reported post-flight reductions in intrinsic connectivity within the right insula and ventral posterior cingulate cortex as well as decreased FC between the cerebellum and motor cortex. It is noteworthy that the cosmonaut exhibited decreases in FC involving the insula and posterior cingulate cortex -- brain areas part of distributed networks that exhibited FC decreases during our HDBR+CO_2_ intervention. Furthermore, we found that this functional encompassing bilateral insular cortices and right PCC exhibited a distinct pattern of FC changes during HDBR+CO_2_ compared to our HDBR control group. This finding suggests that the observed connectivity changes were not due to strict HDBR alone, but rather were due to the combination of strict HDBR and elevated CO_2_.

When comparing our results across resting-state fMRI studies of spaceflight and spaceflight analog environments, the most notable commonalities are FC decreases involving the insula and posterior cingulate cortex. The insula is an integral part of the vestibular network. The vestibular system uses multisensory cues (e.g., visual, somatosensory, auditory, and vestibular cues) for spatial orientation, self-motion perception, gaze stabilization, and postural control (Balaban, 2016). During spaceflight and in spaceflight analog environments, these multimodal cues provide incongruent information, requiring the central nervous system to adaptively reweight sensory inputs. The observed FC decreases among motor, somatosensory, vestibular, cognitive, and multimodal integration brain areas during HDBR+CO_2_ may reflect such multisensory reweighting. The posterior cingulate cortex (PCC) is a core region of the default mode network, which is activated during internally-focused cognition such as introspection, autobiographical memory retrieval, etc. (Buckner et al., 2008). Regions within the PCC are also part of other resting-state networks including the sensorimotor, salience, fronto-parietal, and dorsal attention networks. The functional role(s) of the PCC remain debated, however, theories suggest its involvement in internally-directed thought and controlling balance between external and internal attention (Leech and Sharp, 2014). FC changes involving the PCC may be related to changes in internally-focused cognition during bed rest. These commonalities support the validity of HDBR as a model for studying spaceflight-associated functional neuroplasticity.

Moreover, unique to our HDBR+CO_2_ intervention were FC changes involving visual brain areas, namely primary visual cortex (V1) and lateral occipital cortex (LOC). V1 is involved in processing low-level visual features (Tootell et al., 1998). LOC is involved in visual object recognition, but is especially active when viewing inverted visual stimuli (e.g., faces and objects) (Aguirre et al., 1999). This is an interesting finding since the subjects’ heads were tilted down and they would have had a partially inverted view of the experimenters and their surroundings. Although a non-significant trend compared to our HDBR control group, our observation of reduced ICC within V1 during HDBR+CO_2_ is interesting in light of a recent ‘multihit’ hypothesis that elevated CO_2_ levels onboard the ISS contribute to the development of ophthalmic abnormalities and visual impairments collectively known as Spaceflight Associated Neuro-ocular Syndrome (SANS)(Zwart et al., 2017; Smith and Zwart, 2018), which affects approximately 30% of astronauts returning from long-duration spaceflight (Mader et al., 2011; Mader and Robert Gibson, 2017). This multihit hypothesis posits that a cascade of anatomical and physiologic changes in microgravity (e.g., CO_2_-induced cerebral blood flow increases, headward fluid shifts, vascular leakage, edema) block CSF drainage which increases pressure on the optic nerve and eye, ultimately resulting in neuro-ocular impairment (Zwart et al., 2017; Smith and Zwart, 2018). It is feasible that increases in pressure exerted on the posterior segment of the eye and/or the optic nerve could affect visual inputs to V1 and contribute to reduced intrinsic connectivity. However, future study is needed to investigate this issue. Interestingly, Laurie et al. (2019) collected optical coherence tomographic images before and at the end of HDBR+CO_2_. They found that 5 of the 11 subjects in the current study developed optic disc edema --one of the signs of SANS -- during the intervention. To our knowledge, this is the first HDBR study to observe ophthalmic changes in participants. Comparisons of brain connectivity changes between the subgroups who did and did not develop signs of SANS during HDBR+CO_2_ will be reported in a subsequent paper.

We also investigated if FC changes following HDBR+CO_2_ were associated with behavioral performance alterations across subjects. We used a rod and frame test to estimate subjects’ visual perception of true vertical. On average, subjects’ responses became more consistent (i.e., lower response variability) during the HDBR+CO_2_ intervention. This may reflect practice effects and/or the use of sensory reweighting as a strategy to improve trial-to-trial response consistency. We found that FC decreases between visual, vestibular and somatosensory brain areas were associated with increases in response consistency on the rod and frame test following bed rest. FC reductions between brain areas within the default mode network (i.e., PCC, SFG, and SPL (Buckner et al., 2008)) was also associated with improvements in response consistency. In contrast, subjects who exhibited increased FC between default mode network areas or between visual, vestibular and somatosensory brain areas tended to show increased response variability. The observed functional decoupling between visual, somatosensory and vestibular-related brain areas may reflect changes in sensory weighting that ultimately improve visual orientation perception following HDBR+CO_2_.

Subjects exhibited varying degrees of slowing on the Functional Mobility Test following our HDBR+CO_2_ intervention. FC increases between primary motor cortex (M1) and V5 (also known as hMT+, a higher order visual region involved in visual motion processing (Zeki et al., 1991)) and between premotor cortex and Crus I of the cerebellum were associated with greater slowing on the Functional Mobility Test after bed rest. Those subjects who exhibited decreases in FC between these areas showed less slowing. This finding suggests a link between functional decoupling among motor and higher order visual areas and less deterioration of locomotor abilities following HDBR+CO_2_. We also found that FC decreases between V1 and inferior parietal lobule (IPL) were associated with greater slowing on the Functional Mobility Test. Subjects who exhibited FC increases between V1 and IPL showed less deterioration on the Functional Mobility Test. Primary visual cortex is involved in low-level visual processing (Tootell et al., 1998). It is believed that the IPL is involved in performing sensorimotor transformations (Rushworth et al., 1998), with some studies also reporting its involvement during preparation of foot movements (Sahyoun et al., 2004), and learning sequences of foot movements (Lafleur et al., 2002). This result suggests a compensatory benefit of V1 and IPL interactions following HDBR+CO_2_.

Prolonged exposure to elevated CO_2_ levels may affect brain vascular function and neural activity, which cannot be disassociated using fMRI (Buxton, 2013). In the current study, Laurie et al. (2019) acquired measures of end-tidal PCO_2_, which did not show a significant change from the beginning to the end of the HDBR+CO_2_ intervention. Laurie et al. (2020) also recently reported that arterial partial pressure of carbon dioxide (PaCO_2_) and cerebrovascular reactivity to CO_2_ did not differ between the baseline and HDBR+CO_2_ phases. However, analyses of blood samples collected as part of NASA’s standard measures assessment in the current study showed a small but significant increase in PaCO_2_ from pre-to post-HDBR+CO_2_. Changes in cerebral perfusion may therefore contribute to the observed brain connectivity changes. However, since we did not have an ambulatory + CO_2_ group, we cannot fully determine if any of the observed brain changes are due to elevated CO_2_ exposure alone.

A limitation of the current study is the small sample sizes for the HDBR+CO_2_ (n=11) and HDBR control (n=8) groups. While the inclusion of FC data from the HDBR control group provides insight into whether the observed FC changes are due to strict HDBR or the combined effects of CO_2_ during strict HDBR, the HDBR control group is not ideal. First, the HDBR control data were collected as part of a subsequent bed rest study which was conducted approximately one year following data collection for the HDBR+CO_2_ group. The timeline of the HDBR control group differs from that of the HDBR+CO_2_ group, with the HDBR control group undergoing 60 days of HDBR. The HDBR control group also included few females (6 males, 2 females). Finally, practice effects on the rod and frame test performance present a further limitation. Response variability on the rod and frame test showed a clear learning curve across sessions. While post-bed rest increases in response variability support an effect of the HDBR+CO_2_ intervention, it is currently not possible to disentangle effects of practice from the effects of the HDBR+CO_2_ intervention.

Here we investigated the time courses of resting-state FC changes from before, during, to after 30 days of HDBR in elevated CO_2_. Our hypothesis-driven approach showed FC increases among visual, vestibular, and motor brain areas during HDBR+CO_2_ which decreased following bed rest as well as widespread FC decreases among vestibular, visual, somatosensory and motor brain areas which increased following bed rest. Our hypothesis-free approach revealed a trend whereby FC decreases within primary visual cortex during HDBR+CO_2_. We further examined relationships between FC changes and behavioral changes associated with HDBR+CO_2_. We found that post-bed rest FC changes between visual, vestibular, and somatosensory regions were associated with changes in the consistency of visual perception of true vertical. Finally, we found that post-bed rest FC changes between visual and motor brain areas related to slowing of locomotor performance across subjects. We propose that these alterations of resting-state functional connectivity are a reflection of bed rest-associated multisensory reweighting and CO_2_-related changes in cerebral perfusion. Our findings provide novel insight into the functional neuroplastic mechanisms underlying adaptation to HDBR+CO_2_ microgravity analog environment. This knowledge will further improve HDBR as a model of microgravity exposure, and contribute to our knowledge of brain and performance changes during and after spaceflight.

